# A change in the cell wall status initiates the elimination of the nucellus in Arabidopsis

**DOI:** 10.1101/2024.04.23.590775

**Authors:** Wenjia Xu, Dennys-Marcela Gomez-Paez, Sandrine Choinard, Miryam Iannaccone, Elisa Maricchiolo, Alexis Peaucelle, Aline Voxeur, Kalina T Haas, Andrea Pompa, Enrico Magnani

## Abstract

The evolution of the seed habit coincides with a change in the cell fate of the nucellus, the sporophytic tissue responsible for female meiosis. Seeds arose when the nucellus retained the female spores instead of releasing them in the environment. As a consequence, the nucellus was partially eliminated to accommodate the growth of the female gametophyte inside the sporophyte. With the evolution of angiosperm seeds, the process of nucellus elimination was requisitioned to allow the growth of the endosperm, the second fertilization product devoted to store nutrients. Cell elimination differs from most known cell death programs as it leads to the apparent dismantling of the cell wall. Here, we show that nucellus elimination in Arabidopsis is initiated by the lysis of the pectic polysaccharides in the cell wall. This process exposes other cell wall components to possible further degradation and precedes a cell death program that leads to nuclear DNA fragmentation. Both pathways are regulated by TRANSPARENT TESTA 16, a MADS-domain transcription factor that evolved with seed plants. Finally, the causal effect of cell wall modification on nucellus development is demonstrated by inhibiting pectin degradation, thus suggesting that a change in the cell wall status might have driven seed evolution.

## INTRODUCTION

The plant sexual reproduction cycle-meiosis, sex differentiation, and fertilization-times the alternation of gametophytic and sporophytic generations (*1*). Mitosis intercalates the cycle phases and determines the predominance of one generation versus the other. Finally, the physical separation of gametophytic or sporophytic generations guarantees the dispersal of the progeny.

In heterosporous non-seed plants, the nucellus sporophytic tissue (also referred to as megasporangium) undergoes meiosis and releases female spores in the environment, thus separating the female gametophyte from the sporophyte (*2*). The evolution of the seed habit is marked, instead, by the retention of the female spores in the nucellus and the dispersal of the zygotic embryo, the next sporophytic generation (*3, 4*). Seed plants evolved therefore the ability to eliminate part of the nucellus tissue in order to accommodate the growth of the female gametophyte inside the sporophyte. With the evolution of angiosperm seeds, the nucellus is further consumed by the endosperm, the second fertilization product responsible for storing nutrients and nourishing the embryo (*4–7*). Such changes in the cell fate of the nucellus allowed a ‘revolution’ in human society, as seeds are the foundation of agriculture and human diet.

The signaling pathway leading to the elimination of the nucellus has been elucidated in Arabidopsis. A first round of cell elimination is initiated in the ovule during the growth of the female gametophyte and is marked by a maximum auxin response (*8–10*). After fertilization, the endosperm triggers the elimination of a few more cell layers of the nucellus (hereafter referred to as “transient nucellus”, TN) (Figure 1A and B) (*11*). The AGAMOUS LIKE 62 (AGL62) MADS-domain transcription factor is supposed to send a signal from the endosperm to relieve the repressive action mediated by FERTILIZATION INDEPENDENT SEED (FIS) Polycomb group (PcG) proteins on TN elimination. Downstream of FIS PcG proteins, the TRANSPARENT TESTA 16 (TT16) MADS-domain transcription factor promotes the elimination of the TN. Finally, a few proximal cell layers of the nucellus (hereafter referred to as “persistent nucellus”, PN) (Figure 1A and B) survive throughout seed development and play a role in sugar transport (*12*).

**Figure 1.**
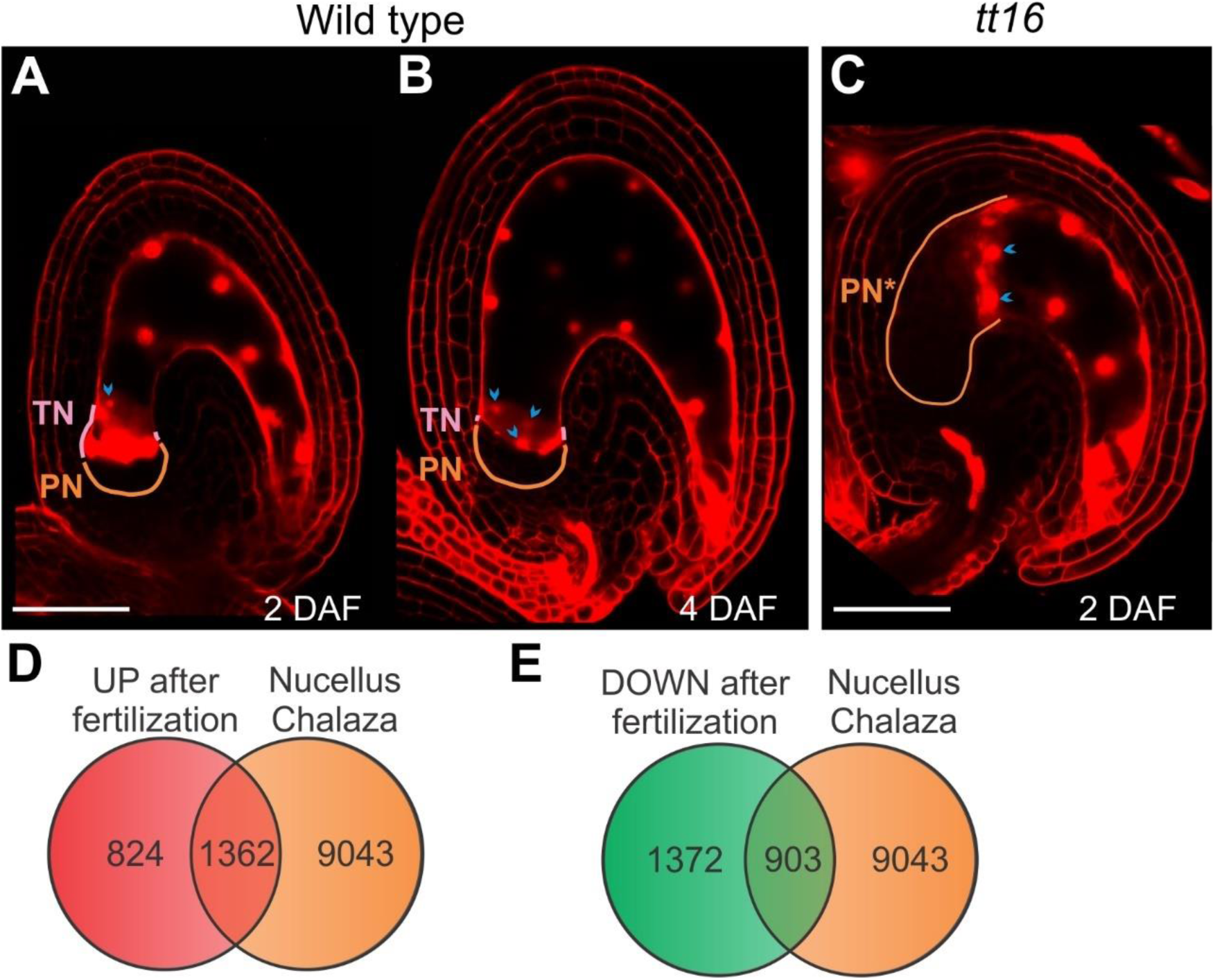
Transcriptional changes induced by fertilization in the chalaza/nucellus tissue. **(A-C)** Propidium iodide fluorescence images of whole mount wild type and *tt16* seeds. DAF, days after flowering; TN, transient nucellus (purple); PN, persistent nucellus (orange); PN*, PN+TN not eliminated (orange). Cyan arrowheads indicate endosperm nuclei close to the nucellus. Bar = 50 µm. **(D)** and **(E)** Venn diagram of genes expressed in the chalaza-nucellus tissue and up-regulated or down-regulated across fertilization.

Compared to most other cell death programs, which preserve the cell wall, cell elimination is characterized by the complete dismantling of unwanted cells, leaving no easily detectable cell corpse (*13*). Images of the TN, obtained by transmission electron microscopy, displayed cell walls consumed to the breaking point and the release of intracellular material in the apoplastic space (*11*). Nevertheless, the process of cell wall elimination in the TN has never been investigated beyond its morphological characterization. Primary cell walls are mostly composed of structural proteins and three families of polymers: cellulose, hemicelluloses, and pectins (*14, 15*). Cellulose accounts for 30–40% of the wall mass and constitutes the main backbone of the cell wall. Hemicelluloses are a diverse family of polymers, including xyloglucans, xylans, and mannans, that strengthen the cell wall by interacting with cellulose. Finally, pectins are heterogeneous polysaccharides rich in galacturonic acid accounting for up to 35% of primary cell walls. To characterize how the cell walls of the TN are eliminated, we conducted an *in silico* analysis of transcriptomic data across fertilization, which pointed toward a change in pectin metabolism. A combination of immunolabeling, genetic and biochemical analyses showed that the pectic polysaccharide homogalacturonan (HG) is de-methylesterified and eventually degraded by pectate lyases in the TN, at the onset of cell elimination. Furthermore, histochemical studies revealed that HG degradation exposes cellulose to possible further degradation. We discovered that HG de-methylesterification is promptly followed by DNA fragmentation, a hallmark of cell death, and that both processes are promoted by TT16. Finally, we demonstrated that inhibition of HG de-methylesterification arrests TN elimination, thus indicating that modulation of the pectin status influences nucellus cell fate and might have played a key role in seed evolution.

## RESULTS

### Fertilization induces the expression of pectin modifier genes in the nucellus

In Arabidopsis, the elimination of the transient nucellus (TN, Figure 1A and B) is characterized by the apparent clearance of the entire cell corpses including the cell walls (*11*). To investigate this unique process, we conducted a transcriptomic analysis aimed at identifying cell wall-related genes expressed in the nucellus in response to fertilization. We sorted genes differentially expressed (DEGs) across fertilization (log2 fold change equal or greater than one) in transcriptomic data generated by Figueiredo and colleagues (Fig. 1D and E and Table S1) (*16*). Among these DEGs, we selected genes expressed in the nucellus/chalaza tissue of seeds at the pre-globular embryo stage, according to transcriptomic data obtained by laser capture microdissection (Fig. 1D and E and Table S1) (*17*). Finally, we performed a Gene Ontology (GO) analysis and looked for GO terms related to the cell wall. Among genes up-regulated after fertilization and expressed in the chalaza-nucellus tissue, we detected significant enrichments in GO annotations for “pectin metabolic process” (p-value 7.17E-04) and “cell wall modification” (p-value 9.76E-04) (Table S1). In particular, *PECTIN METHYLESTERASEs* (*PMEs*) and *PECTATE LYASEs* (*PLs*) were highly represented (Table 1). It has been shown that homogalacturonan (HG), the most abundant pectin subtype in plant cell walls, is synthetized as methylesterified polymer (*18*). De-methylesterification of HG by PMEs can lead to its degradation by PLs and influence cell wall integrity (*19*), thus being a primary suspect in the elimination of the nucellus and therefore the focus of our investigation.

**Table 1.**
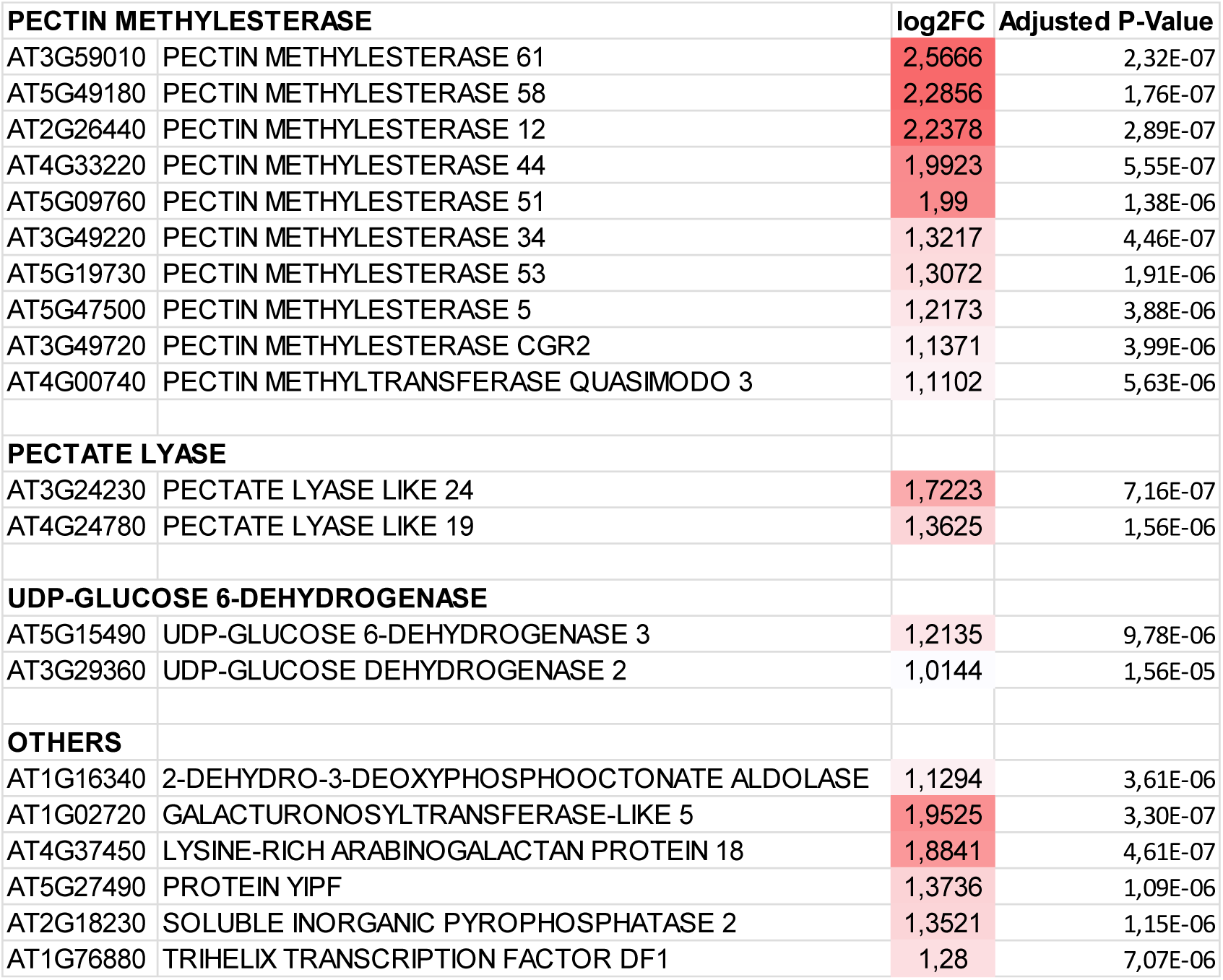
Pectin-related genes transcriptionally regulated by fertilization in the chalaza/nucellus tissue. Genes annotated with the GO term “pectin metabolic process” and up-regulated after fertilization in the chalaza-nucellus tissue. Log2 fold changes in expression across fertilization and adjusted P-Values, as obtained by Figueiredo and colleagues, are annotated on the right (*16*).

### Pectin de-methylesterification and degradation mark the onset of nucellus elimination

To test if HG in the TN cell walls is de-methylesterified and degraded, we stained whole mount seeds with propidium iodide (PI), a dye that binds HG carboxyl residues and nucleic acids (Figure 1A and B) (*20*). PI fluorescence in the cell wall has been shown to increase in the presence of PMEs, which create more binding sites, and decrease in the presence of PLs, which digest HGs (*20*). To identify the nucellus, we used the inner integument 1, the pigment strand and the endosperm as morphological markers, as previously described (*11*). PI fluorescence was visible as amorphous signal in the TN at 2 days after flowering (DAF), during the process of cell elimination, and almost completely disappeared at 4 DAF when the TN was degraded (Figure 1A and B). PI also stained endosperm nuclei, recognizable by their round shape (Figure 1A and B). By contrast, we detected a relatively lower PI signal in the PN at both time points tested (Figure 1A and B). These data suggest that HG is de-methylesterified (at 2 DAF) and degraded (at 4 DAF) in the TN whereas it remains highly methylesterified in the PN.

To further test this hypothesis, we conducted immunolabeling studies using 2F4 and LM20 antibodies, which are specific for low and high methylesterified pectins, respectively (*21, 22*). During the first 4 DAF, we detected a higher 2F4 signal in the TN nucellus, when compared to the PN (Figure 2A-C and Figure S1). This was true even at 1 DAF (Figure 2A), before any morphological marker of nucellus elimination is visible. To precisely analyze the pattern of HG methylesterification, we quantified the fluorescent signal of the 2F4 and LM20 antibodies at 2 DAF. We observed a decrease in the relative HG methylesterification status along the proximal-distal axis of the nucellus (Figure 2E and F, see materials and methods). Overall, these results indicate that HG de-methylesterification is a hallmark of the TN.

**Figure 2.**
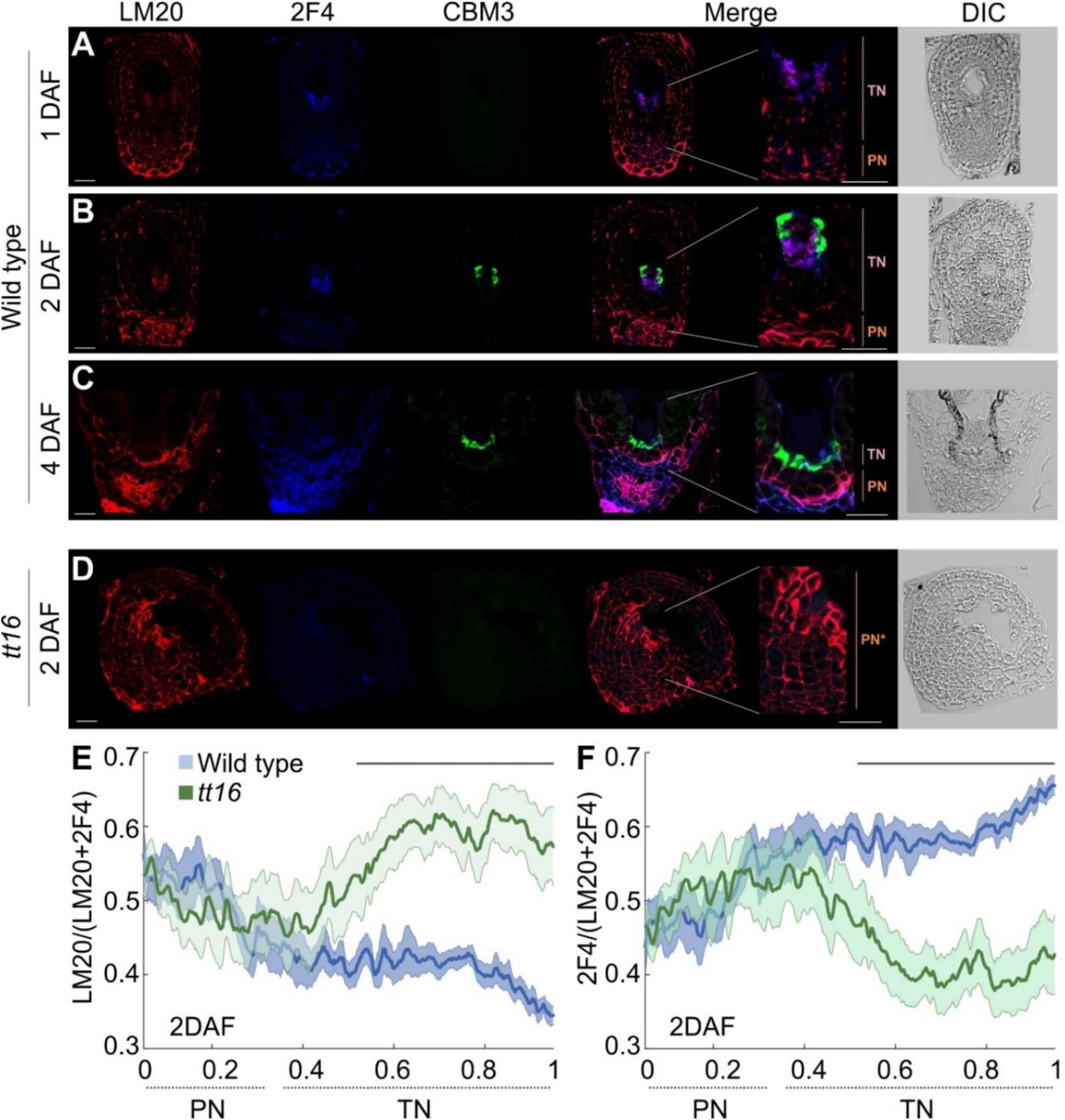
Pectin methylesterification status in the nucellus. **(A-D)** Triple labeling of wild type and *tt16* seed sections with LM20 (in red) and 2F4 (in blue) antibodies and CBM3 (in green) probe. DIC, differential interference contrast image; DAF, days after flowering; TN, transient nucellus (purple); PN, persistent nucellus (orange); PN*, PN+TN not eliminated (orange). Bars = 20 µm. **(E)** and **(F)** Mean relative fluorescent signal of LM20 and 2F4 antibodies along the wild type and *tt16* nucellus proximal-distal axis (see materials and methods). Lines at the top of the graph indicate regions of statistically significant difference between wild type and *tt16* (Wilcoxon test, *P* < 0.05). Error bars indicate standard error.

To determine if de-methylesterified HG in the TN is targeted by pectin-degrading enzymes, we tested our samples with the carbohydrate binding module 3 (CBM3) probe, which binds crystalline cellulose in cells actively depositing cellulose or characterized by a relatively low abundance of pectins (*23*). The cellulose sites bound by CBM3 are indeed masked by pectins, thus making CBM3 an inverse pectin marker (*23*). In support of the hypothesis of pectin degradation, we detected CBM3 fluorescence in the TN of seeds at 2 and 4 DAF, in proximity to 2F4 signal (Figure 2B and C and Figure S1). By contrast, the TN of seeds at 1 DAF displayed 2F4 but not CBM3 fluorescence suggesting that HG de-methylesterification precedes its digestion (Figure 2A). Furthermore, the PN did not show CBM3 fluorescence at any developmental stage tested (Figure 2A-C).

To confirm if pectins are digested in the TN, we characterized the localization of the PECTATE LYASE LIKE 24 (PLL24) enzyme, which we detected in our *in silico* analysis (Table 1). In line with transcriptomic data obtained by laser capture microdissection (*17*), the *PLL24* promoter and genomic region drove expression of the GREEN FLUORESCENT PROTEIN (*ProPLL24:PLL24-GFP*) in the TN of seeds at 2 and 4 DAF (Figure 3A and B). This result suggests that the HG degrading machinery is present during TN elimination.

**Figure 3.**
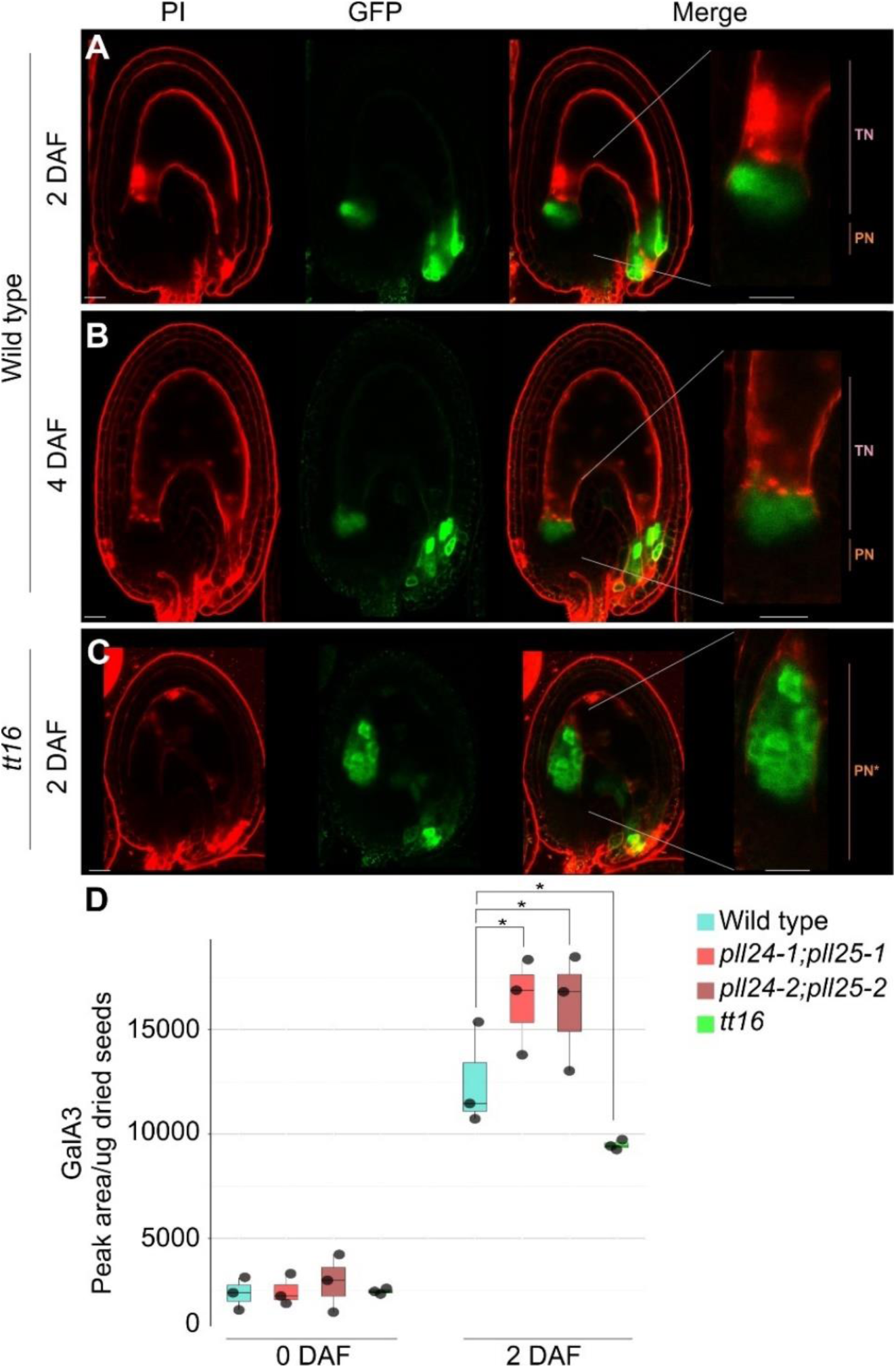
Pectin degradation in the nucellus. **(A-C)** Propidium iodide (PI, in red) and GFP (in green) fluorescence images of whole mount *ProPLL24:PLL24-GFP* wild type and *tt16* seeds. TN, transient nucellus (purple); PN, persistent nucellus (orange); PN*, PN+TN not eliminated (orange). Bar = 20 µm. **(D)** Profiling of the 3-residue galacturonic acid oligomer (GalA3) in wild type, *pll* and *tt16* ovules and seeds. Asterisks indicate statistically significant difference between wild type and mutant lines (Student’s t-test, P<0.05). Error bars indicate standard deviation. DAF, days after flowering.

To challenge our model, we compared the amount of digestible HG present in wild type versus two independent double *pll24;pll25* mutant lines. *PLL25* is the closest paralogue of *PLL24* (*24*) and its expression is up-regulated after fertilization (Table S1) in the chalaza-nucellus tissue at the embryo globular stage (*17*). To this end, we quantified the amount of HG oligomers resulting from the digestion of wild type and *pll24;25* mutant cell walls with an endo-polygalacturonase, which hydrolyzes low and non-methylesterified HGs, using a mass spectrometry sensitive analytical method based on the separation of oligosaccharides combined with accurate determination of their sizes and methylesterification patterns (*25, 26*). In ovules at 0 DAF, we did not detect any statically significant difference between wild type and *pll24;25* mutant samples (Figure 3D). By contrast, digestion of *pll24;pll25* seeds at 2 DAF released significantly more 3-residue galacturonic acid oligomer (GalA3), indicating a higher content of digestible HG compared to the wild type (Figure 3D). In line with our previous analyses, these results indicate that PLL24 and PLL25 actively digest low-methylesterified HG in response to fertilization.

### Nuclear DNA fragmentation follows pectin de-methylesterification in the transient nucellus

To determine if a cell death program works alongside the breakdown of the cell wall and partially pre-digests the intracellular material, we tested the TN for DNA fragmentation, a hallmark of programmed cell death, by terminal deoxynucleotidyl transferase dUTP nick end labeling (TUNEL) (*27, 28*). We started to observe TUNEL signal at 2 DAF in the most distal cells of the TN (Figure 4A and B), when compared to positive and negative controls (Figure S2). The signal disappeared in seeds at 4 DAF, where only the PN was visible (Figure 4C). These results demonstrate that nuclear DNA fragmentation follows HG de-methylesterification and works alongside HG degradation toward the complete elimination of the TN.

**Figure 4.**
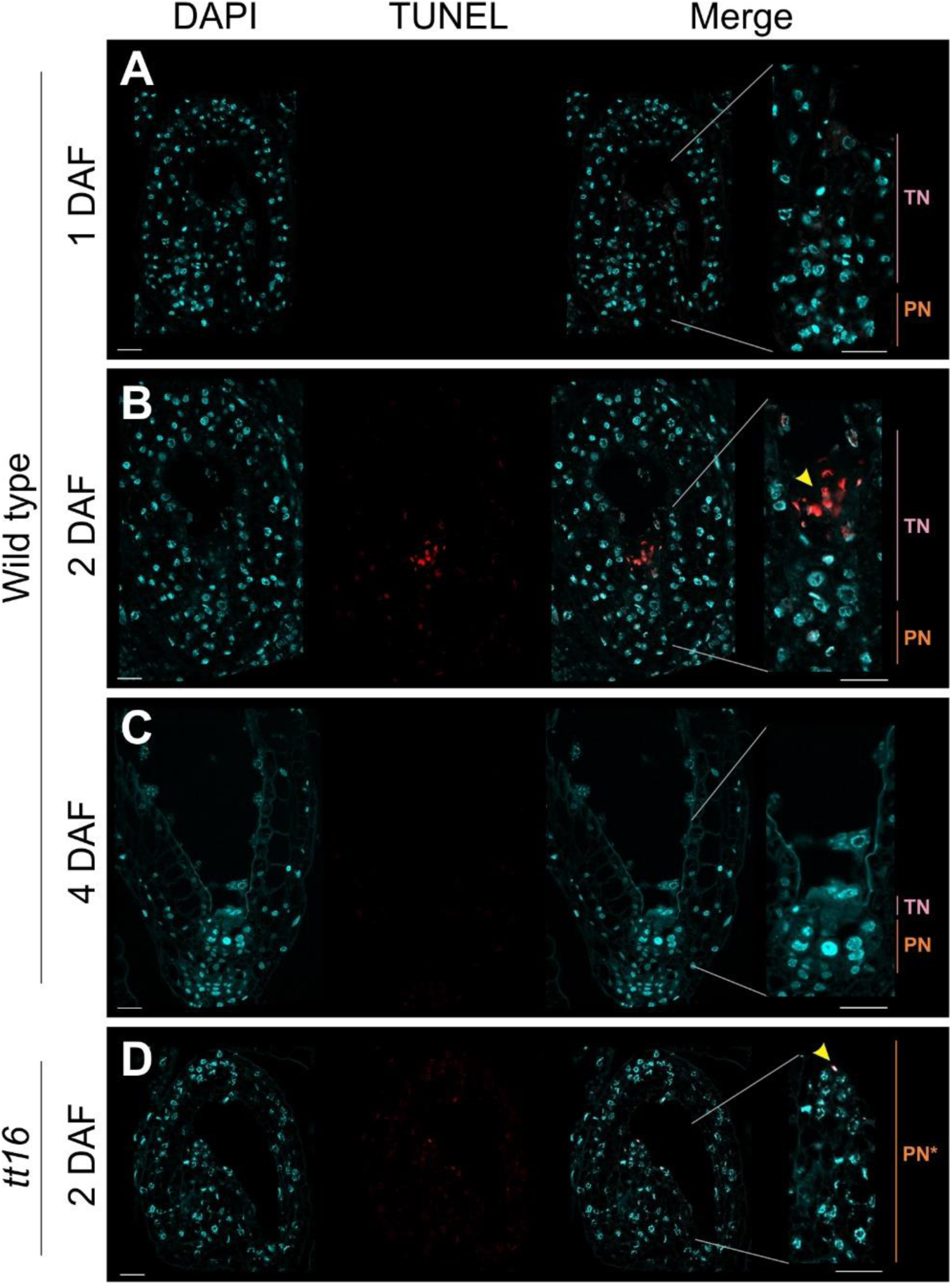
Nuclear DNA fragmentation in the nucellus. **(A-D)** DAPI (in cyan) and TUNEL (in red) fluorescence images of wild type and *tt16* seed sections. Yellow arrowheads indicate TUNEL positive nuclei. DAF, days after flowering; TN, transient nucellus (purple); PN, persistent nucellus (orange); PN*, PN+TN not eliminated (orange). Bar = 20 µm.

### TT16 promotes pectin de-methylesterification and nuclear DNA degradation in the transient nucellus

The process of nucellus elimination is promoted by the TT16 transcription factor (*11*). We therefore challenged our model in *tt16* mutant seeds, which display an almost intact nucellus (referred to as PN* because it is made of PN and TN not eliminated) where only a few distal cells degenerate (Figure 1C). In contrast to what observed in the wild type, the entire nucellus of *tt16* seeds at 2 DAF displayed relatively high-methylesterified HG both in propidium iodide and immunolabeling experiments (Figure 1C and 2D-F). Furthermore, we did not detect CBM3 signal in the nucellus indicating that HG is not degraded in the mutant (Figure 2D). Nevertheless, *PLL24* expression was not affected by the *tt16* mutation (Figure 3C), a result that supports the knowledge that HG degradation requires its de-methylesterification. In line with this hypothesis, mass spectrometry analyses detected a lower content of the GalA3 oligomer, and therefore digestible HG, in *tt16* mutant seeds treated with endo-polygalacturonase, when compared to the wild type (Figure 3D). Overall, these data suggest that TT16 promotes cell wall elimination in the TN by inducing HG de-methylesterification.

Finally, we addressed if both HG modification and DNA fragmentation are regulated by TT16. To this end, we conducted TUNEL experiments and detected a faint signal in only a few distal cells of the *tt16* nucellus at 2 DAF (Figure 4D), when compared to the wild type (Figure 4B). Altogether, our results indicate that TT16 regulates HG de-methylesterification and nuclear DNA degradation in a coordinated fashion to achieve full elimination of the TN.

### Pectin de-methylesterification is necessary for nucellus elimination

To test if HG de-methylesterification is necessary to trigger cell elimination, we induced the over-expression of the *PECTIN METHYL ESTERASE INHIBITOR 3* (*PMEI3*) gene, which is known to inhibit PME activity and block HG de-methylesterification in Arabidopsis (*29, 30*). To this end, we employed an ethanol-inducible system based on the constitutive *35S* promoter (*iPMEI3-OX*) and over-expressed the *β-glucuronidase* (*GUS*) gene as control *(iGUS-OX*) (*29*). At 4 DAF, 70% of *iPMEI3-OX* seeds displayed an enlarged and correctly developed endosperm (Figure 5B and S3) but did not undergo TN elimination (Figure 5B), similar to *tt16* seeds (Figure 5D) (*11*). By contrast, the TN of all *iGUS-OX* seeds analyzed was almost completely eliminated (Figure 5A) as in the wild-type (Figure 5C). These data demonstrate that HG de-methylesterification is necessary for the elimination of the TN.

**Figure 5.**
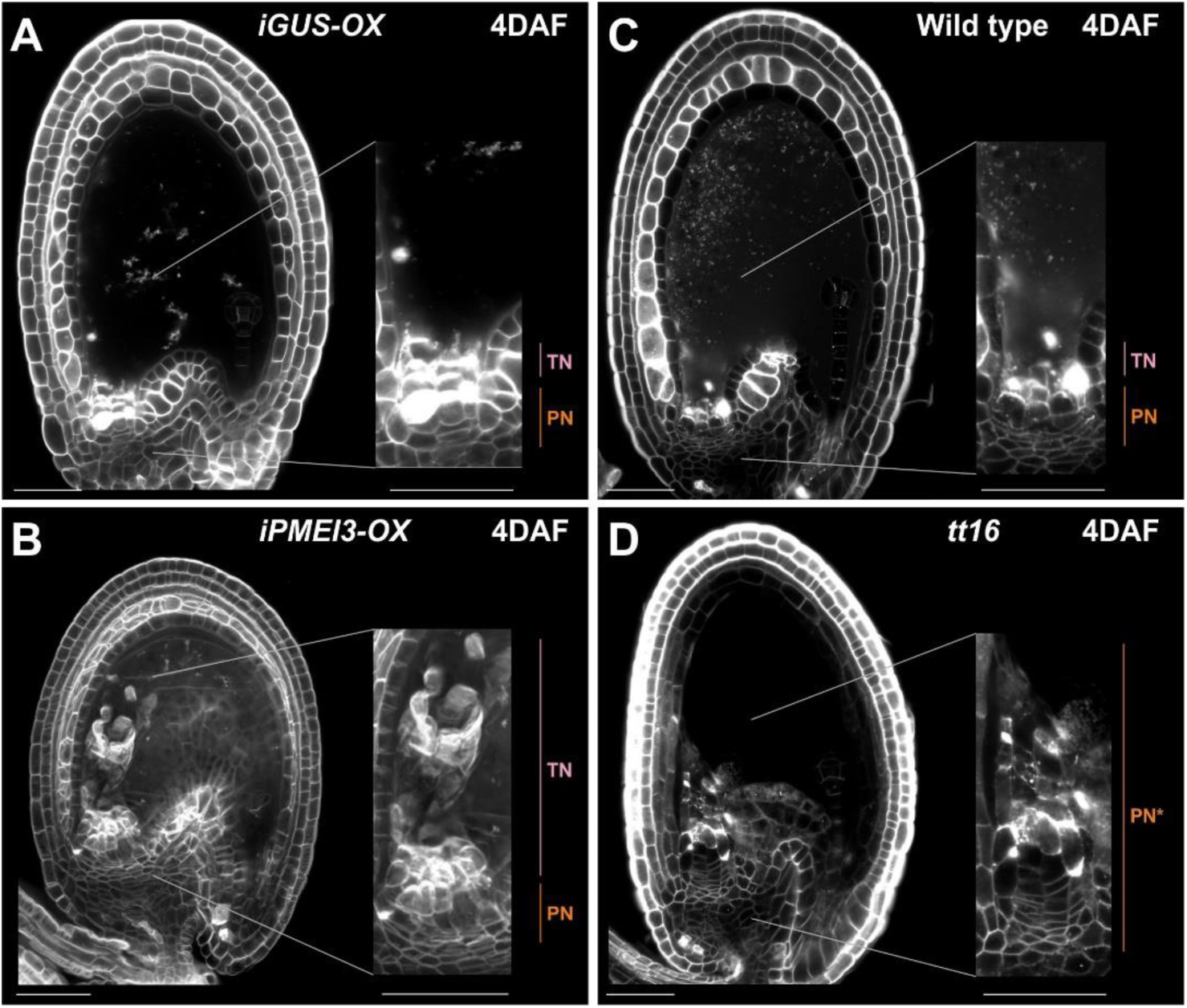
Inhibition of pectin de-methylesterification arrests nucellus elimination. **(A and B)** Renaissance 2200 fluorescence images of *iGUS-OX*, *iPMEI3-OX*, wild type and *tt16* seeds. DAF, days after flowering; TN, transient nucellus (purple); PN, persistent nucellus (orange); PN*, PN+TN not eliminated (orange). Bar = 50 µm.

## DISCUSSION

Nucellus elimination was instrumental to seed evolution as it allowed the growth of female gametophyte and endosperm inside the maternal sporophyte. Compared to most other cell death programs, cell elimination appears to dismantle the cell wall, a unique process that was not fully addressed at the molecular level. Here, we show that de-methylesterification of the cell wall homogalacturonan (HG) triggers the elimination of the nucellus, thus suggesting extracellular polysaccharides had a key role in the evolution of the seed.

### Dismantling the cell wall starting from its pectic component

The cell wall is a unique component of plant cells that provides strength, rigidity, and protection, through its chemical composition that is hard to break down (*14, 15*). Indeed, most cell death programs do not eliminate the cell wall but only the intracellular content (*13, 27*). The nucellus tissue, with few other examples (*13*), constitutes an exception to such a rule as its most distal cell walls are dismantled to reorganize the internal space of the seed and make room for the growth of the endosperm. This process had been already described in a number of species, including Arabidopsis, but only at the morphological level (*5*). Here, we identify pectin degradation as the gate to nucellus elimination by analyzing transcriptomic data obtained across fertilization. Our analysis revealed, indeed, a significant enrichment in the GO term “pectin metabolic process” among genes differentially expressed right after fertilization. In particular, we detected ten *PMEs* and two *PLLs* genes as up-regulated after fertilization. The list of functions attributed to pectins has been expanding in recent years. Originally considered important for intercellular adhesion because of their abundance in the middle lamella, pectins have been now shown to be involved in cell fate specification, morphogenesis, intercellular communication, and environmental sensing (*31*). HG is secreted into the cell wall as methylesterified polymer (*18*) and then subjected to de-methylesterification via the action of wall-bound pectin methylesterases (PMEs), which ultimately affect the mechanical properties of the cell wall (*19*). PMEs produce free carboxyl groups that can interact with Ca2+ to create a pectic gel, thus increasing cell wall stiffness. Alternatively, low methylesterification of HG is recognized as cleavage site by PECTIN/PECTATE LYASE (PL) or POLYGALACTURONASE (PG) enzymes that trigger HG de-polymerization, thereby promoting cell wall loosening. The rapid transcriptional up-regulation of the pectin degrading machinery observed after fertilization in the TN suggested that the cell walls are deprived of their pectic component to proceed further into cell elimination. Immunolabeling analyses confirmed that HG is preferentially low methylesterified in the TN after fertilization, when clear morphological signs of nucellus elimination are not yet visible. HG de-methylesterification is therefore the first known hallmark of nucellus elimination and is rapidly followed by the detection of crystalline cellulose. We hypothesize that cellulose is accessible only after HG degradation in non-dividing or expanding cells. This scenario was confirmed by detecting PLL24 in the TN during its elimination. Furthermore, we showed that the amount of degradable HG was higher in two independent *pll24/25* mutant lines, when compared to the wild type. These data suggest that PLL24 and PLL25 actively degrade HG in the TN, exposing other cell wall polymers, such cellulose, to possible further degradation.

Modification of the pectin status in the TN is regulated by the TT16 transcription factor, which promotes cell elimination. *tt16* seeds displayed preferentially high methylesterified HG in the distal nucellus, a pattern diametrically opposed to what observed in the wild type. By contrast, *PLL24* expression was not affected by the *tt16* mutation. We therefore suggest that TT16 promotes TN elimination by inducing HG de-methylesterification, the first step toward pectin degradation. Similarly, the PN of wild type seeds showed a higher level of HG methylesterification, when compared to the TN, which might guarantee its survival. In support of this hypothesis, the inhibition of HG de-methylesterification by *PMEI3* over-expression arrested the elimination of the TN. This result clearly shows a causal effect of cell wall modification on the development of the nucellus. Our model relates to studies conducted in cereals. In barley ovules, HG in the distal nucellus has been shown to be low methylesterified and interpreted as responsible for keeping the cells more rigid and less prone to degradation (*32*). In light of what presented here, we believe that this hypothesis should be revised in favor of a role of HG de-methylesterification in initiating cell wall degradation. Consistent with our results, Qin Sun *et al.* showed that the maize EXPANSIN B15, a protein responsible for cell wall loosening, promotes the process of nucellus elimination (*33*). We speculate that expansins might facilitate the work of PME and PLL enzymes by relaxing the cell wall. Both articles describe an effect of nucellus development on grain size, thus opening the way to novel approaches for yield-improvement. Our results might contribute to the design of lines showing a faster or slower elimination of the nucellus by regulating the HG esterification status.

### Evolving a seed by modifying the cell wall

The elimination of the nucellus marked the evolution of the seed habit as it allowed the growth of the spores inside the sporophyte instead of releasing them. The same mechanism was exploited by angiosperms to evolve endospermic seeds, such as Arabidopsis and cereals, which favor the growth of the endosperm at the expense of the nucellus. Here we show that the process of pectin de-methylesterification is necessary to trigger the dismantling of the nucellus cell walls. Therefore, we speculate that a change in the pectin status might have started such evolutionary breakthroughs that had a profound impact on our society.

A similar set of events has been described during pathogen attacks. The first step in the infection of plant cells by necrotrophic fungi is indeed the de-methylesterification and degradation of pectins to improve cell wall accessibility to other degrading enzymes and achieve a successful penetration (*34*). Another parallel can be drawn between the elimination of the nucellus and the softening of the endosperm during embryo growth. In Arabidopsis, the bHLH-type transcription factor ZHOUPI (ZOU), similarly to TT16, promotes weakening of the endosperm cell walls by regulating the activity of a series of cell wall-modifying enzymes (*35, 36*). In *zou* mutant seeds, the (1–5)-α-L-arabinan epitopes, which might be associated with rhamnogalacturonan-1 pectins, were more abundant and persistent than in the wild type. These data suggest that ZOU might soften the endosperm cell wall by modifying pectins. Finally, the clearance of dying cells in animals is undertaken by phagocytes, which engulf and digest cell corpse and extracellular matrix (*37*). Despite their profound differences, the plant cell wall and the animal extracellular matrix are both composed of polysaccharides and proteins (*38*). Similar to HG in plants, hyaluronic acid (HA) polymers are non-protein polysaccharides within the mammalian extracellular matrix with pivotal structural and signaling roles in programmed cell death, cellular mobility, and the regulation of cancer (*39*). A number of studies has associated changes in the molecular weight of HA polymers with apoptosis and phagocytic elimination in cancer cells (*39*). The striking convergence in both the structure and function of HA and HG emphasizes the crucial contribution of sugar-derived polymers to the emergence of multicellular organisms.

### Dying while the cell wall is eliminated

The elimination of the cell wall might lead to the death of the TN nucellus cells without invoking any intracellular cell death program. Such a scenario seemed to be suggested by transmitted electron microscope images showing cell corpses released in the apoplast during nucellus elimination (*11*). Nevertheless, we detected DNA fragmentation in the nuclei of TN cells after HG de-methylesterification but before cell wall dismantling. Nuclear DNA degradation has been detected in a number of developmental cell death programs (*27, 40*), thus suggesting that the TN undergoes programmed cell death as well as cell wall degradation. Combining both processes might allow the TN to pre-digest cellular material before the collapse of the cell walls. Further processing of the cell corpses might then continue in the apoplast.

TUNEL experiments in *tt16* seeds showed that only a few distal cells of the TN nucellus undergo DNA fragmentation while most of the nucellus expand in apparent coordination with the rest of the maternal tissues. These data suggest that PCD and cell wall degradation are either independent processes co-regulated by TT16 or inter-dependent pathways downstream of TT16. During the elimination of the endosperm cells, ZOU promotes weakening of the cell walls while NAC transcription factors induce PCD (*35, 36, 41*). In the *zou* mutant, the endosperm displays intact cell walls but expression of *NAC* and PCD marker genes, despite being delayed. These data indicate that ZOU transcription factor is principally responsible for the softening of the cell wall and not PCD. TT16 and ZOU might therefore regulate cell elimination in a different fashion. Alternatively, PCD in *tt16* seeds might be substantially delayed. Further experiments are necessary to identify the genes responsible for PCD in the nucellus to test such hypotheses.

## MATERIALS AND METHODS

### Genetic material

The *tt16-1* allele was isolated in the Wassilewskija accession from the INRA Versailles collection (*42*) and then backcrossed to Columbia-0 (*43, 44*). *pll24-1* (SALK_049124), *pll24-2* (SALK_ 035767), *pll25-1* (SALK_031335), and *pll25-2* (SALK_068375) are in the Columbia-0 accession. *iPMEI3-OX* and *iGUS-OX* are in the in the Wassilewskija accession (*45*).

### Accession Numbers

*TT16* (*AT5G23260*), *PLL24* (*AT3G24230*), *PLL25* (*AT4G13710*), and *PMEI3*

(*AT5G20740*).

### Molecular biology

*PLL24* 4.6 kb promoter and genomic region was PCR amplified using the gene-specific primers (*PLL24F 5′-AGCGTGACGGATTTGGATTGA-3′* and *PLL24R 5’-ACGGCAACTAAGTGCACCTGC-3’*) carrying the *attB1* (*5′-GGGGACAAGTTTGTACAAAAAAGCAGGCT-3′*) and *attB2* (*5′-GGGGACCACTTTGTACAAGAAAGCTGGGTC-3′*) Gateway recombination sites at the 5′-ends, respectively. The PCR product was amplified with the high-fidelity Phusion DNA polymerase (Thermo Fisher Scientific), recombined into the *pDONR221* vector (BP Gateway reaction) according to the manufacturer’s instructions (Thermo Fisher Scientific), sequenced, and then recombined into the *pMDC107* binary vector (*46*).

### GO enrichment analysis

GO enrichment analyses were conducted on the GENEONTHOLOGY website using the PANTHER classification system (*47*).

### Mass spectrometry analysis

Seeds (600 for each sample, n = 3) were submerged in 96% (v/v) ethanol and ground. The pellets were collected by centrifugation (13,000 g for 10 min) and dried in a speed vacuum concentrator at 30°C overnight. Samples were digested with 1 U/mg dried weight of Pectobacterium carotovorum endo-Polygalacturonanase (Megazyme, Bray, Ireland) in 50 mM ammonium acetate buffer (pH 5) at 37°C for 18 h. After digestion, samples were centrifuged at 13,000 rpm for 10 min, and 100 µL of the supernatants were transferred into vials. For mass spectrometry analysis, 10 μL of each fraction was injected into the machine. The oligosaccharides released from digestion were separated according to Voxeur et al., 2019 (*25*). Chromatographic separation was performed on an ACQUITY UPLC Protein BEH SEC Column (125Å, 1.7 μm, 4.6 mm × 300 mm, Waters Corporation, Milford, MA, USA) coupled with a guard Column BEH SEC Column (125Å, 1.7 μm, 4.6 mm × 30 mm). Elution was performed in 50 mM ammonium formate, 0.1% formic acid at a flow rate of 0.4 mL/min, with a column oven temperature of 40°C. The injection volume was set to 10 μL. Quantitative evaluation of cell wall fragments was conducted using an HPLC system (UltiMate 3000 RS HPLC system, Thermo Scientific, Waltham, MA, USA) coupled to an Impact II Ultra-High Resolution Qq-Time-Of-Flight (UHR-QqTOF) spectrometer (Bruker Daltonics, Bremen, Germany) equipped with an electrospray ionization (ESI) source in negative mode. The end plate offset set voltage was 500 V, capillary voltage was 4000 V, nebulizer pressure was 40 psi, dry gas flow was 8 L/min, and the dry temperature was set to 180°C. The Compass 1.8 software (Bruker Daltonics) was used to acquire the data, and peak areas were integrated manually.

### Ethanol-inducible gene expression system

*iPMEI3-OX* and *iGUS-OX* plants were treated with ethanol as previously described (*29*). Nevertheless, ethanol induction of *PMEi3* tended to arrest seed development and we therefore analyzed siliques of untreated plants, which displayed a milder *PMEI3* over-expression phenotype due to the leakiness of the system. We imaged 46 *iPMEI3-OX* and 32 *iGUS-OX* seeds.

### Microscopy

Wild type and GFP-expressing whole mount seeds were analyzed 15 min after mounting in a 100 µg·mL-1 propidium iodide, 7% sucrose solution (Weight/Volume).

For staining with Renaissance 2200, seeds were immersed in a NaOH (0.2 M), SDS (1%) solution at 37 °C for 3 h, washed three times in water, transferred to a fresh bleach solution (2%) for 10 min to remove precipitated tannins, washed five times, and then mounted in Renaissance 2200 solution diluted 100 times with water.

Immunolabeling and CBM3 histochemical analyses were conducted as previously described (*23*). Wild type and mutant samples at the same developmental stage were mounted on the same slide.

For TUNEL experiments we used the In Situ Cell Death Detection Kit, TMR red (Roche) and followed the manufacturer’s instructions.

Samples were imaged by confocal laser scanning microscopy (Leica SP8 and Zeiss LSM 710) keeping conditions uniform between samples to allow cross-comparisons.

### Quantitative analysis of immunolabeling experiments

The multisegment linear region of interest was drawn manually using Fiji with a line thickness spanning the entire nucellus. Subsequent analyses were carried out in Matlab, utilizing custom-written scripts available on GitHub. The raw data comprised the total intensity (summed over the line thickness) for two imaging channels at specific points along the line segment. The imaged seeds exhibited variability in size due to natural differences in developmental stages. To consolidate the data, we assumed that seeds preserved the proportion between persistent and transient nucellus regardless of their size. Consequently, we interpolated the data using the same number of query points for all seeds, employing Matlab’s built-in function interp1, and the interpolation grid was generated using the linspace function. Finally, the data were normalized as follows: IN-LM20 = ILM20/(ILM20 + I2F4), IN-2F4 = I2F4/(ILM20 + I2F4), where ILM20 and I2F4 represent the staining intensity of methylated and demethylated homogalacturonans, respectively. IN-LM20 and IN-2F4 are presented as averages (n = 10), ± the standard error of the mean. Comparison of IN-LM20 and IN-2F4 data between wild type and *tt16* was performed at each query point using the Wilcoxon rank test at a confidence level of 0.05.

## AUTHORS CONTRIBUTIONS

W.X. performed the research and analyzed the data. D.M.G.P. performed part of the immunolabeling analyses. S.C. cloned *PLL24*, identified the *pll* mutations and helped to perform genetic analyses. M.I., El.M. and An.P. characterized *PLL24* expression. A.P. helped to perform immunolabeling analyses, analyze the data and write the article. A.V. performed mass spectrometry analyses. K.T.H. quantified immunolabeling results. E.M. designed the research, analyzed the data and wrote the article.

## ACKNOWLEDGEMENTS

We thank Gwyneth Ingram, Herman Höfte, Samantha Vernhettes and members of their groups for helpful discussions and the Observatoire du Végétal for plant culture, access to imaging and mass spectrometry facilities and assistance.

## LIST OF SUPPLEMENTARY MATERIALS

Figs. S1 to S3 Table S1

## REFERENCES

1. S. N. Bai, The concept of the sexual reproduction cycle and its evolutionary significance. Front Plant Sci 6, 11 (2015).

2. J. A. Banks, Selaginella and 400 million years of separation. Annu Rev Plant Biol 60, 223–238 (2009).

3. J. Pettitt, Heterospory and the origin of the seed habit. Biol. Rev. 45, 401–415 (1970).

4. E. Magnani, Seed Evolution, A’Simpler’ Story. Trends Plant Sci 23, 654–656 (2018).

5. J. Lu, E. Magnani, Seed tissue and nutrient partitioning, a case for the nucellus. Plant Reprod 31, 309–317 (2018).

6. S. Nayar, R. Sharma, A. K. Tyagi, S. Kapoor, Functional delineation of rice MADS29 reveals its role in embryo and endosperm development by affecting hormone homeostasis. J Exp Bot 64, 4239–4253 (2013).

7. L. L. Yin, H. W. Xue, The MADS29 transcription factor regulates the degradation of the nucellus and the nucellar projection during rice seed development. Plant Cell 24, 1049–1065 (2012).

8. J. Wang et al., Auxin efflux controls orderly nucellar degeneration and expansion of the female gametophyte in Arabidopsis. New Phytol 230, 2261–2274 (2021).

9. W. C. Yang, D. Ye, J. Xu, V. Sundaresan, The SPOROCYTELESS gene of Arabidopsis is required for initiation of sporogenesis and encodes a novel nuclear protein. Genes Dev 13, 2108–2117 (1999).

10. U. Schiefthaler et al., Molecular analysis of NOZZLE, a gene involved in pattern formation and early sporogenesis during sex organ development in Arabidopsis thaliana. Proc Natl Acad Sci U S A 96, 11664–11669 (1999).

11. W. Xu et al., Endosperm and Nucellus Develop Antagonistically in Arabidopsis Seeds. The Plant cell 28, 1343–1360 (2016).

12. J. Lu et al., The nucellus: between cell elimination and sugar transport. Plant Physiol 185, 478–490 (2021).

13. G. C. Ingram, Dying to live: cell elimination as a developmental strategy in angiosperm seeds. J Exp Bot 68, 785–796 (2017).

14. Z. A. Popper et al., Evolution and diversity of plant cell walls: from algae to flowering plants. Annu Rev Plant Biol 62, 567–590 (2011).

15. F. SC. (Longman Scientific & Technical, Harlow, 1988), pp. 103–185.

16. D. D. Figueiredo, R. A. Batista, P. J. Roszak, L. Hennig, C. Kohler, Auxin production in the endosperm drives seed coat development in Arabidopsis. Elife 5, (2016).

17. M. F. Belmonte et al., Comprehensive developmental profiles of gene activity in regions and subregions of the Arabidopsis seed. Proc Natl Acad Sci U S A 110, E435–444 (2013).

18. D. Mohnen, Pectin structure and biosynthesis. Curr Opin Plant Biol 11, 266–277 (2008).

19. D. Qiu, S. Xu, Y. Wang, M. Zhou, L. Hong, Primary Cell Wall Modifying Proteins Regulate Wall Mechanics to Steer Plant Morphogenesis. Front Plant Sci 12, 751372 (2021).

20. C. M. Rounds, E. Lubeck, P. K. Hepler, L. J. Winship, Propidium iodide competes with Ca(2+) to label pectin in pollen tubes and Arabidopsis root hairs. Plant Physiol 157, 175–187 (2011).

21. F. Liners, J. J. Letesson, C. Didembourg, P. Van Cutsem, Monoclonal Antibodies against Pectin: Recognition of a Conformation Induced by Calcium. Plant Physiol 91, 1419–1424 (1989).

22. Y. Verhertbruggen, S. E. Marcus, A. Haeger, J. J. Ordaz-Ortiz, J. P. Knox, An extended set of monoclonal antibodies to pectic homogalacturonan. Carbohydr Res 344, 1858–1862 (2009).

23. K. T. Haas, M. Rivière, R. Wightman, A. Peaucelle, Multitarget Immunohistochemistry for Confocal and Super-resolution Imaging of Plant Cell Wall Polysaccharides. Bio Protoc 10, e3783 (2020).

24. L. Sun, S. van Nocker, Analysis of promoter activity of members of the PECTATE LYASE-LIKE (PLL) gene family in cell separation in Arabidopsis. BMC Plant Biol 10, 152 (2010).

25. A. Voxeur et al., Oligogalacturonide production upon. Proc Natl Acad Sci U S A 116, 19743–19752 (2019).

26. A. Paterlini et al., Enzymatic fingerprinting reveals specific xyloglucan and pectin signatures in the cell wall purified with primary plasmodesmata. Front Plant Sci 13, 1020506 (2022).

27. T. Van Hautegem, A. J. Waters, J. Goodrich, M. K. Nowack, Only in dying, life: programmed cell death during plant development. Trends Plant Sci 20, 102–113 (2015).

28. Y. Gavrieli, Y. Sherman, S. A. Ben-Sasson, Identification of programmed cell death in situ via specific labeling of nuclear DNA fragmentation. J Cell Biol 119, 493–501 (1992).

29. K. T. Haas, R. Wightman, E. M. Meyerowitz, A. Peaucelle, Pectin homogalacturonan nanofilament expansion drives morphogenesis in plant epidermal cells. Science 367, 1003–1007 (2020).

30. F. Xu et al., Biochemical characterization of Pectin Methylesterase Inhibitor 3 from. Cell Surf 8, 100080 (2022).

31. Y. Shin, A. Chane, M. Jung, Y. Lee, Recent Advances in Understanding the Roles of Pectin as an Active Participant in Plant Signaling Networks. Plants (Basel) 10, (2021).

32. X. Yang et al., Ovule cell wall composition is a maternal determinant of grain size in barley. New Phytol 237, 2136–2147 (2023).

33. Q. Sun et al., A NAC-EXPANSIN module enhances maize kernel size by controlling nucellus elimination. Nat Commun 13, 5708 (2022).

34. C. P. Kubicek, T. L. Starr, N. L. Glass, Plant cell wall-degrading enzymes and their secretion in plant-pathogenic fungi. Annu Rev Phytopathol 52, 427–451 (2014).

35. S. Yang et al., The endosperm-specific ZHOUPI gene of Arabidopsis thaliana regulates endosperm breakdown and embryonic epidermal development. Development 135, 3501–3509 (2008).

36. C. Fourquin et al., Mechanical stress mediated by both endosperm softening and embryo growth underlies endosperm elimination in Arabidopsis seeds. Development 143, 3300–3305 (2016).

37. A. M. Fond, K. S. Ravichandran, Clearance of Dying Cells by Phagocytes: Mechanisms and Implications for Disease Pathogenesis. Adv Exp Med Biol 930, 25–49 (2016).

38. L. Stavolone, V. Lionetti, Extracellular Matrix in Plants and Animals: Hooks and Locks for Viruses. Front Microbiol 8, 1760 (2017).

39. M. Bhattacharyya, H. Jariyal, A. Srivastava, Hyaluronic acid: More than a carrier, having an overpowering extracellular and intracellular impact on cancer. Carbohydr Polym 317, 121081 (2023).

40. D. Latrasse, M. Benhamed, C. Bergounioux, C. Raynaud, M. Delarue, Plant programmed cell death from a chromatin point of view. J Exp Bot 67, 5887–5900 (2016).

41. N. M. Doll et al., Endosperm cell death promoted by NAC transcription factors facilitates embryo invasion in Arabidopsis. Curr Biol 33, 3785–3795 e3786 (2023).

42. N. Bechtold, G. Pelletier, In planta Agrobacterium-mediated transformation of adult Arabidopsis thaliana plants by vacuum infiltration. Methods Mol Biol 82, 259–266 (1998).

43. O. Coen et al., Developmental patterning of the sub-epidermal integument cell layer in Arabidopsis seeds. Development 144, 1490–1497 (2017).

44. N. Nesi et al., The TRANSPARENT TESTA16 locus encodes the ARABIDOPSIS BSISTER MADS domain protein and is required for proper development and pigmentation of the seed coat. Plant Cell 14, 2463–2479 (2002).

45. A. Peaucelle, R. Wightman, K. T. Haas, Multicolor 3D-dSTORM Reveals Native-State Ultrastructure of Polysaccharides’ Network during Plant Cell Wall Assembly. iScience 23, 101862 (2020).

46. M. D. Curtis, U. Grossniklaus, A gateway cloning vector set for high-throughput functional analysis of genes in planta. Plant Physiol 133, 462–469 (2003).

47. H. Mi, A. Muruganujan, J. T. Casagrande, P. D. Thomas, Large-scale gene function analysis with the PANTHER classification system. Nat Protoc 8, 1551–1566 (2013).

